# Development of an organotypic brain slice culture model using tissues collected from geriatric mice

**DOI:** 10.64898/2025.12.23.696302

**Authors:** Addison Keely, Jackson Wezeman, Savanna Roberts, Sonali Bhana, Warren Ladiges

## Abstract

Ex vivo organ slice cultures are attractive models to address specific biomedical research questions because they retain complex and dynamic three dimensional architecture. In addition, they require only a few animals as tissue donors thus greatly decreasing the number needed for live animal studies. The model has been most successful using tissues from very young mice, but has generally not been considered for aging research because of the challenge of maintaining viability of tissues collected from older mice. The brain is a high impact organ in aging and age-related disease research, so it was of interest to investigate the viability and related molecular and cellular characteristics of brain slice cultures (BSCs) derived from old mice. Coronal sections of 200 μm thickness were collected from the middle third of the brain of 21 month old C57BL/6JN mice, cultured in a standard media concoction for seven days, exposed to chemical stressors glucose, hydrogen peroxide, lipopolysaccharide, or sodium hypochlorite for 24 to 48 hours, and then rinsed and cultured for another seven days. Results in non-treated BSCs showed cellular viability of greater than 80 percent with specific aging pathways relatively unaffected. Chemical stressors selectively targeted pathways of aging including DNA damage, autophagy, and inflammation. These preliminary observations suggest that BSCs derived from geriatric mice have potential for research into brain aging and age-related neurodegenerative conditions, and could serve as a prototype for developing organ slice cultures for other organs collected from old mice.

## Introduction

Organ slice cultures are a promising yet under-utilized and under-appreciated model for research on aging. This ex vivo model consists of very thin slices of tissue that continue to function in a cell culture venue, and has potential for testing response to chemical stressors and gerotherapeutic drugs with minimal animals needed (as tissue donors), with low research costs compared to live animal studies. Organ slice cultures retain the complex and dynamic three dimensional architecture allowing cellular interactions to occur [1]. These advantages provide the opportunity to model organ-specific characteristics of aging at the cellular and molecular level.

Of particular interest for neurodegenerative conditions is the ability to study the role of brain aging using a brain slice culture (BSC) model. Brain slice culturing of neonatal brain tissue has been used in numerous studies [2, 3] but there is concern that neonatal tissue may not typify brain aging observed in older mice. Therefore, BSCs derived from geriatric mice were tested to see if molecules involved in pathways of aging [4] were expressed, and if response to chemical stressors involved aging pathways.

## Methods

C57Bl/6JN mice were obtained from the National Institute on Aging’s Aged Rodent Colony and housed in the mouse vivarium at the University of Washington in Seattle, WA. Mice were euthanized at 21 months of age using cervical dislocation as the primary method. The brain was extracted onto a cold plate and the cerebellum and brainstem were removed. The remaining brain was immediately transferred to ice-cold 1x PBS (Gibco), then adhered to a vibratome plate with a drop of superglue and supported by sterile 1% agarose solution for cutting. Coronal slices of 200 μm were cut using a Vibratome 1500 Sectioning System. Collection of slices began after the first 3 mm of brain tissue were removed starting at the olfactory lobe, approximately where the hippocampus starts (Figure 1). Subsequent 200 μm slices were assigned to alternating treatment groups for representative tissue from the hippocampus in each treatment group. A total of 21 coronal slices were collected over a 4.2 mm rostral length of brain. Three slices were placed in formalin to serve as Control Day 0. All remaining slices were placed on a 0.4 μm pore Nunc™ Polycarbonate cell culture insert in a 6-well plate (Thermo Scientific™) containing 1 mL of culture medium (50% MEM/HEPES (Gibco), 25% heat-inactivated horse serum (Gibco/Lifetech, Austria), 25% Hanks’ solution (Gibco), 2 mM NaHCO3 (Merck, Austria), 6.5 mg/ml glucose (Merck, Germany), 2 mM glutamine (Merck, Germany), pH 7.2, and 1% penicillin/streptomycin (Thermo Scientific™).

**Figure 1.**
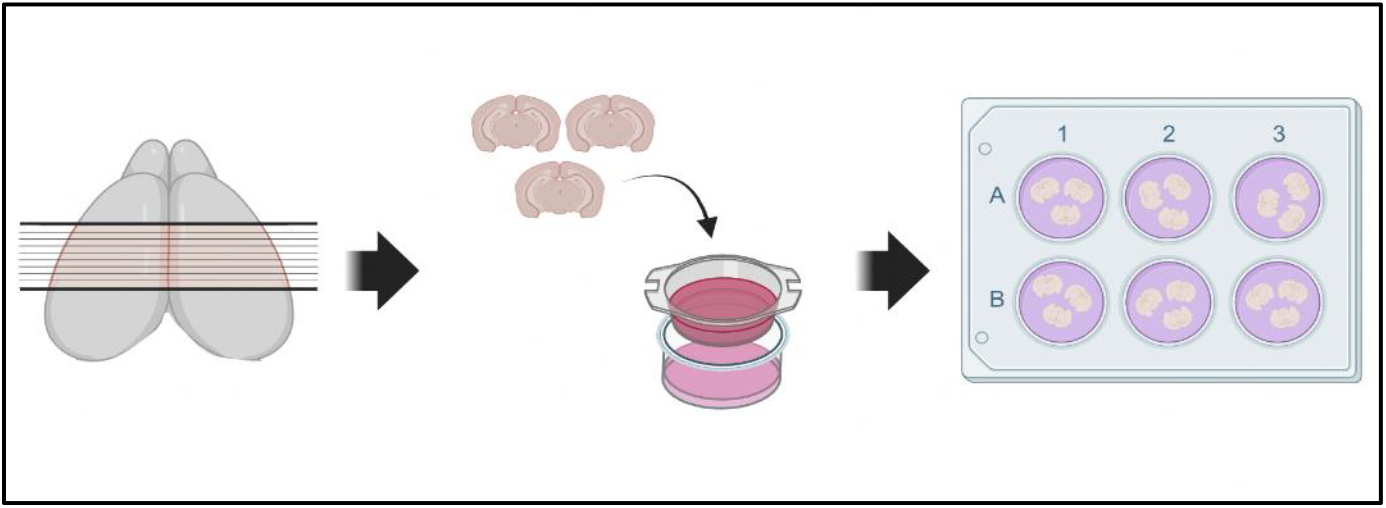
The prepared mouse brain was sliced into 200 μm thick coronal sections with a Vibratome. Collection of 21 total slices (4.2 mm rostral length) began after 3 mm of brain tissue had been removed. Sections were placed on the semi-permeable membrane of a removable cell culture insert in sequential wells of a 6-well plate with 1mL of slice medium. Created in https://BioRender.com

Brain slices were incubated at 37°C and culture medium was changed every 2 to 3 days. One slice well was reserved as an untreated control, labelled Control day 14. After seven days in culture, chemical stressors were added to all remaining wells: hydrogen peroxide (ThermoScientific Cat. No: 426000010) at 150 μM for 48 hours [5], glucose (Fisher Chemical D-Glucose Anhydrous D16-3) at 25 mM for 48 hours [6], lipopolysaccharide (Sigma Aldrich L4391) at 50ng/mL for 48 hours (unpublished lab data), and sodium hypochlorite (Chlorox) at 5 μM for 24 hours [7]. Media with chemical stressors were then removed and wells rinsed twice with 1x PBS before adding regular media and resuming culture conditions for the remaining seven days.

After 14 days, brain slices were transferred to cassettes and fixed in 10% neutral buffered formalin for 24 hours before being transferred to PBS for storage. Slices were embedded in type 6 paraffin wax (Fisher Scientific) according to standard embedding procedures. Histology slides were created from blocks using a microtome at 4 μm thickness on a charged glass slide.

Cresyl Violet staining (Abcam ab246817) was used to visualize surviving neurons in the slices, based on stained Nissl substances in the endoplasmic reticulum. Staining was performed by rehydrating the slides in a 70 minute stepwise series as in IHC staining, then incubating slides in 1% Cresyl Violet stain solution for 5 minutes followed by dehydration in three washes of 100% ethanol then 10 minutes in xylene before adhering coverslips to slides with toluene permount. Slides were allowed to dry for 24 hours before imaging and analysis.

IHC was performed using kits from Abcam (HRP/DAB Rabbit Kit: ab64261, HRP/DAB Mouse Kit: ab64259) according to manufacturer protocols. Slides are rehydrated in a stepwise series running from xylene, 100% ethanol, 95% ethanol, 70% ethanol, and deionized water for a total running time of 70 minutes. Slides were then placed in a hot water bath with a citrate buffer antigen retrieval (pH 6.0, Sigma-Aldrich: C9999) for 20 minutes. Slides were allowed to cool for 10 minutes, and then rinsed in the wash buffer (1x TBST; Tris-buffered saline with Tween20) before a hydrogen peroxide buffer block was applied to the tissue. Slides were washed in the wash buffer for 10 minutes. A protein block applied that reduces non-specific binding of the target antibody, and slides were then washed for 10 minutes as previously described. Antibody was diluted in the wash buffer according to recommended dilution (yH2AX ThermoFisher 10856-1-AP at 1:200 and CCL2/MCP1 Novus NBP1-07035 at 1:800). Slides incubated with the diluted antibody overnight in a 4°C fridge. On the second day, the diluted antibody was washed, then slides incubated with biotinylated goat anti-mouse or anti-rabbit followed by streptavidin peroxidase, with washes after each. DAB was then applied in a fume hood for 2 minutes per slide. Slides were then washed with the wash buffer then deionized water for 15 minutes, then dehydrated in a reverse-series for 5 minutes per solution going from 70% ethanol, 95% ethanol, 100% ethanol, and finally xylene. Coverslips were adhered on the slides using toluene permount (Fisher). Slides were air dried for 24 hours before imaging.

Images were taken of all stained tissue at 20X with a Nikon DS-Fi3 camera on a Nikon Eclipse Ci microscope using NIS Elements software. Images were run through ImageJ Analysis (Mac OS X 10.16) through a QuPath Program (0.3.2) to determine the percent density of positive staining on the tissues. Unpaired two-tailed t-tests were performed to analyze the differences between means of control and treated groups. Shapiro-Wilk tests and Q-Q plots indicated no evidence of non-normality in the data sets. All data analysis was performed with GraphPad Prism version 10.4.2.

## Results and Discussion

Preliminary experiments showed cell viability greater than 90% after 14 days in culture using Trypsin-EDTA (Thermo Scientific) for tissue dissociation and Trypan blue (Thermo Scientific) stain for cell viability. Viability counts were performed using Accuris QuadCount™ Automated Cell Counter. In addition, representative brain slices at 14 days were incubated with Trypan blue for three minutes then rinsed with PBS. Visual counts at 3x magnification showed cell survival at 80-90%. For neuron-specific assessment of viability, cresyl violet staining was used to show that neuronal density decreased by 23 percent indicating some neuronal degeneration and death, but the majority of neurons were still viable after 14 days in culture (Figure 2).

**Figure 2.**
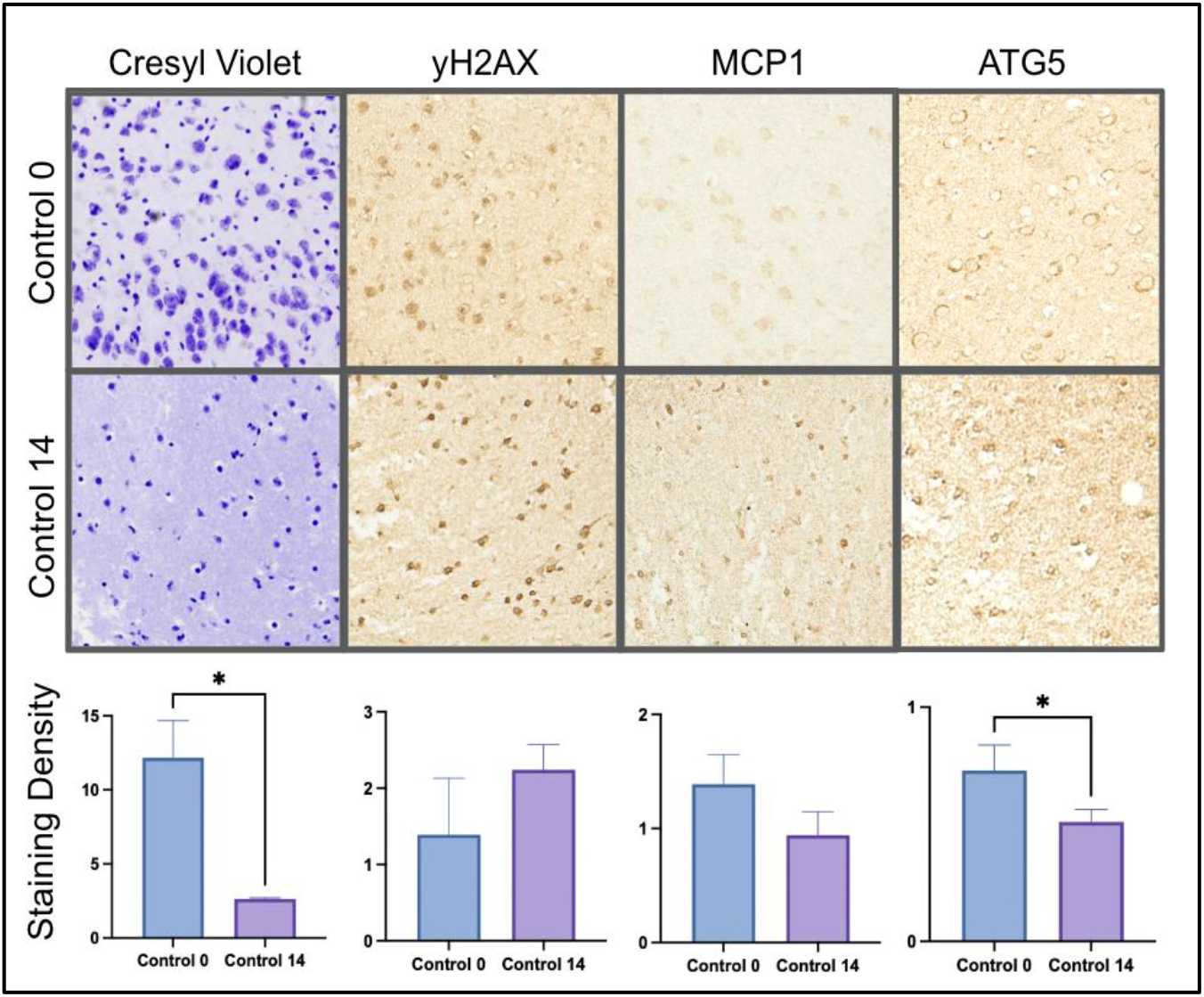
Digital images at 200x magnification showed identification of molecules representing pathways of aging in control day 0 slices compared to control day 14 slices. Cresyl violet staining was significantly lower in cultured BSCs (p<0.05). There was no significant difference in yH2AX staining between groups (p=0.1). MCP1 staining was slightly higher in Control 14 slices (p=0.07). ATG5 staining was significantly higher in control day 0 slices (p<0.05). n=3 for each group.

To see if any changes in expression of molecular markers of aging pathways occurred during the 14 day incubation period, immunohistochemistry and digital imaging were carried out using antibodies optimized for mouse brain tissue. No significant changes occurred for yH2AX (Figure 2) suggesting culture techniques did not cause a significant amount of DNA damage. H2AX is phosphorylated to yH2AX, which triggers a double stranded DNA damage repair response.

Likewise, there was no significant difference in staining intensity for MCP1 (monocyte chemoattractant protein-1), a chemokine used as a biomarker for inflammation. However, IHC staining of ATG5, a protein involved in autophagy, showed an increase in staining intensity in brain slices cultured for 14 days compared to non-cultured brain slices (day 0) suggesting culture conditions had some effect on the autophagy pathway.

It was also of interest to see if recognizable glial cells were present within the brain slices. IHC showed the presence of astrocytes and microglia at Day 0 and after 14 days of culture (Figure 3).

**Figure 3.**
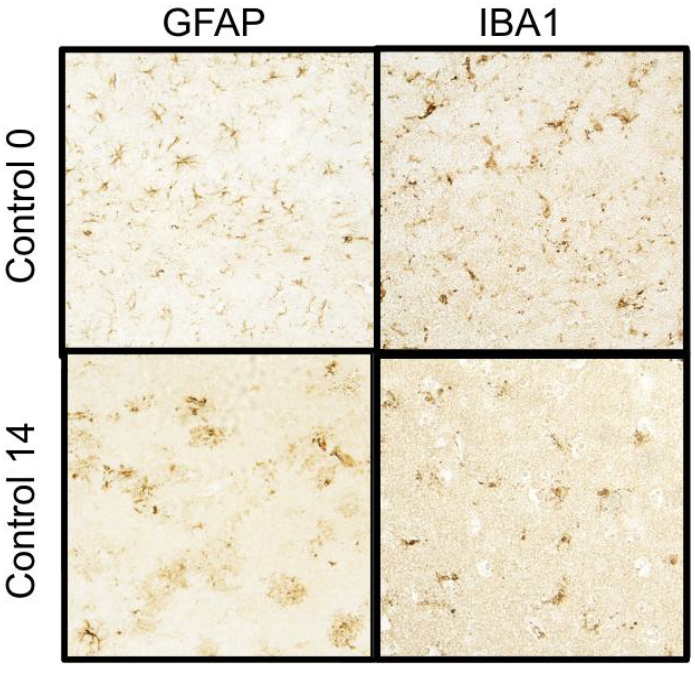
IHC on untreated control BSCs after 14 days in culture show presence of astrocytes and microglia at 400x magnification.

The role of lipopolysaccharide (LPS) in inflammation pathways explains the increased staining intensity of MCP1 in treated BSCs (Figure 4). LPS is an endotoxin that triggers an immune response through toll-like receptor 4 (TLR4) [8], and is an established *in vitro* model of neuroinflammation. LPS also increased staining intensity of ATG5. Autophagy can protect the cell in response to stress [9] and is upregulated during inflammation by a variety of signals [10]. The observations raise the question of why increased autophagy was not sufficient to mitigate the LPS-induced increase in MCP1 expression, as both markers were significantly elevated in the treated group versus the day 14 control.

**Figure 4.**
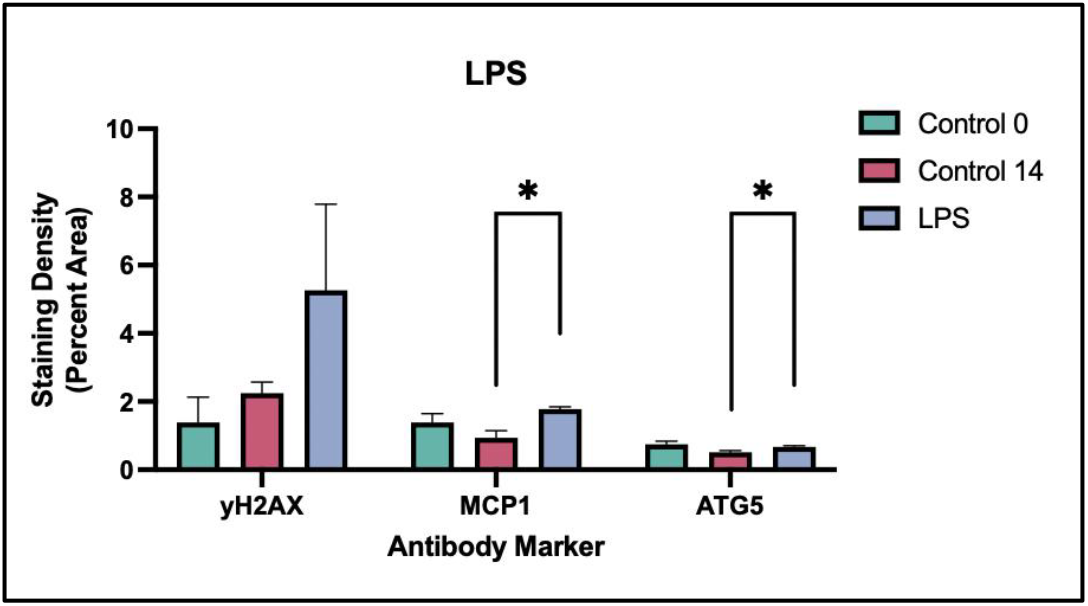
Brain slice cultures treated with LPS had significantly higher inflammation (p<0.05) visualized through MCP1 staining, and autophagy (p<0.05) visualized through ATG5 staining. Differences in DNA damage between LPS and control BSCs were not significant (p=0.07).

BSCs treated with high glucose levels showed a significant increase in ATG5 staining intensity but no differences in staining intensities for yH2AX or MCP-1 (Figure 5). The relationship between high glucose levels and autophagy is complex with multiple involved pathways.

**Figure 5.**
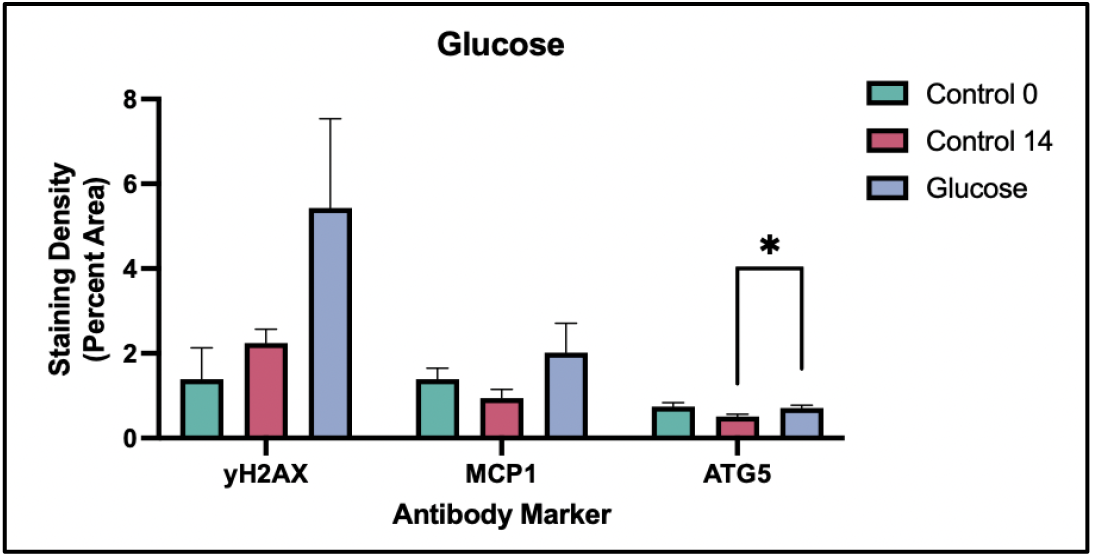
BSCs treated with high glucose had insignificantly higher averages in yH2AX staining for double stranded DNA damage (p=0.06) and MCP1 staining for inflammation (p=0.06). ATG5 staining for autophagy was significantly higher in high glucose-treated BSCs (p<0.05).

Autophagy is upregulated in starvation conditions and suppressed during nutrient abundance to maintain energy homeostasis [11]. The shift from high glucose levels to normal glucose levels, as performed in this study, has previously been shown to induce a state of metabolic stress in microglia, leading to an increase in autophagy [12]. High glucose levels are expected to increase inflammation [13] and DNA breaks [14]. There was a trend for higher average staining of MCP1 and yH2AX though the results were not statistically significant. The glucose concentration used (25 mM) was chosen to optimize cellular stress and minimize cell death, and may have been insufficient to trigger these pathways.

BSCs treated with hydrogen peroxide showed an increased need for DNA repair (Figure 6). Phosphorylated H2AX (yH2AX) has a key role in the DNA repair process [15]. Hydrogen peroxide has been well established as a form of stress in *in vitro* studies to increase ROS presence [16]. Increased ROS levels in the brain cause oxidative stress and neuronal degeneration. The lack of significant differences in MCP1 and ATG5 suggests that the oxidative damage induced by hydrogen peroxide treatment was not severe enough to trigger significant increases in either pathway.

**Figure 6.**
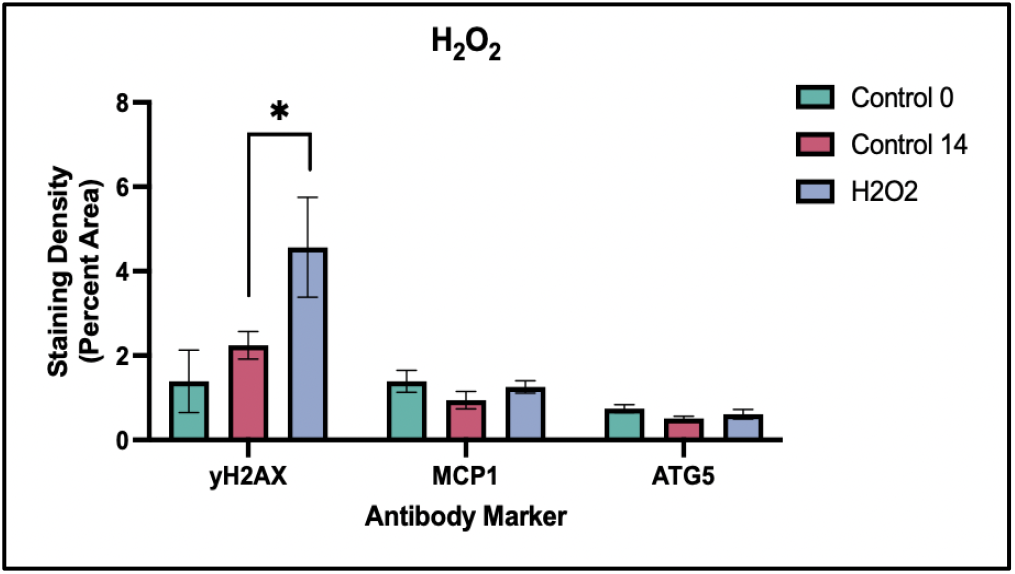
Mouse BSCs treated with hydrogen peroxide had significantly higher staining intensities of yH2AX (p<0.05). Differences in MCP1 and ATG5 staining were not significant (p=0.1 to p=0.2).

Treatment with NaOCl triggered a DNA damage response in BSCs, shown by increased levels of yH2AX (Figure 7), a marker for DNA repair signaling. NaOCl, widely used for its antimicrobial effects, works through oxidation, damaging proteins and other molecules in the cell [17]. The interaction between hypochlorite and DNA bases as well as the denaturation of secondary structure causes DNA damage [18]. The NaOCl dose was low enough to preserve cell functions, as the yH2AX marker was significantly increased in treated BSCs compared to the day 14 controls. This suggests that DNA damage response pathways remained functional and were activated in response to DNA damage in the BSCs. NaOCl treatment also led to increased expression of ATG5. MCP1 was not affected.

**Figure 7.**
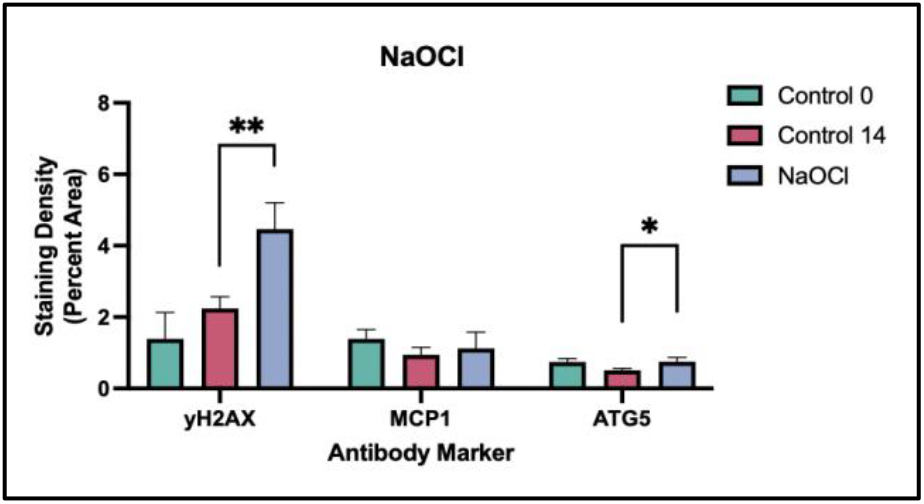
NaOCl treated brain BSCs had significantly higher levels of yH2AX (p<0.01) and ATG5 staining for autophagy (p<0.05) compared to day 14 controls. The differences in inflammation, visualized through MCP1 staining, were not significant (p=0.5).

In conclusion, preliminary evidence suggests that brain slice cultures derived from old mice can retain viability for 14 days under enriched culture conditions. Culture conditions in general supported the ability of cells to express molecules involved in pathways of aging. The BSC model was able to respond to chemical stressors by triggering selected pathways of aging. Observations from this study suggest old-age-mouse derived BSCs have potential for research into brain aging and age-related neurodegenerative conditions, and could serve as a prototype for developing organ slice cultures for other organs collected from geriatric mice.

## Acknowledgements

This study was supported by NIH grant R01 AG067193 (W Ladiges, PI)

